# Losses and gains of fallows impact farmland bird populations over three funding periods of the EU Common Agricultural Policy

**DOI:** 10.1101/2021.10.11.463895

**Authors:** Lionel R. Hertzog, Norbert Röder, Claudia Frank, Hannah G. S. Böhner, Johannes Kamp, Sebastian Klimek

**Author notes:** **correspondence:** Thünen Institute of Biodiversity, Bundesallee 65, 38116 Braunschweig, Germany.

## Abstract

Fallow land provides habitat for threatened and declining farmland biodiversity. Policy change under the EU’s Common Agricultural Policy (CAP) has been driving the area of fallows over the past decades and influenced trends in farmland biodiversity.
We analyzed the relationships between fallow land area across Germany over three CAP funding periods and species richness and abundance of farmland birds. We examined whether the strength of the relationships with fallow land area were moderated by species habitat preferences and landscape configurational complexity (edge density). We combined spatial data on fallow land area from three agricultural censuses in Germany (2007, 2010 and 2016) with country-wide farmland bird monitoring data.
Farmland bird species richness and the abundance of the majority of the studied farmland bird species were positively related to increases in fallows across three CAP funding periods. The relationship of fallows with bird richness was strongest at intermediate levels of edge density. There was generally little support for a moderating effect of edge density on the relations between fallows and bird abundance.
We conclude that the loss of fallows in the period 2007 to 2016 resulted in strong declines of farmland birds. We predict that a future increase of the proportion of fallow land to 4% of the arable land, as envisaged in the German 2023-2027 CAP strategic plans, or to 10%, as foreseen in the EU Biodiversity Strategy, will lead to increases in farmland bird species richness and abundance depending on the landscape context and species-specific habitat preferences.

**Policy implications:** Increasing the proportion of fallow land will likely be a key lever to stabilize and revert negative farmland bird population trends. An increase of fallow area in all but the least complex landscapes will boost farmland bird richness and abundance. Increasing the proportion of fallow land to 4% should bring farmland bird richness and abundance back to the levels observed in 2007 acknowledging that farmland bird populations were already severely depleted in 2007. A more ambitious expansion of fallow land towards 10% should be targeted towards areas that experienced the strongest loss of fallows and towards landscapes with intermediate levels of edge density.

## 1. Introduction

Agricultural intensification is a major global driver of biodiversity loss (IPBES, 2019). Intensification comprises several processes leading to the simplification and homogenization of landscapes through a decline in landscape heterogeneity (e.g. loss of semi-natural habitats, increasing field size), but also an increasing use of agrochemicals, farm mechanization and specialization (Tscharntke et al. 2005, Kleijn et al 2009, Emmerson et al. 2016). As a consequence, biodiversity in agricultural landscapes has been declining across Europe and North America since at least the 1960s (Stanton et al., 2018). In addition, declines in abundance of farmland animals and plants (Meyer et al. 2013), range retractions (Eichenberg et al. 2021), community reorganization and even country-wide species extinctions (Ollerton et al. 2014) following agricultural intensification have been documented.

An important component of intensification is the loss of non-productive features in agricultural landscapes. The availability of non-productive features such as field margins, hedgerows and fallow land plays an important role in driving biodiversity patterns (Van Buskirk & Willi 2004, Sálek et al. 2018). Fallow land in agricultural landscapes provides undisturbed habitat for plants, and food and shelter for animals across trophic levels (Van Buskirk & Willi, 2004, Tscharntke et al., 2011). In Europe, fallows have been part of farming systems since early cultivation (Allen 2000). In the last decades, fallows have become a prominent conservation measure in the European Union (EU) for promoting farmland biodiversity in agricultural landscapes and fallow area is therefore largely driven by supranational and national policies.

In the EU, agricultural land-use patterns are heavily influenced by the Common Agricultural Policy (CAP). The CAP is the main policy instrument used to distribute public payments in the agricultural sector. Its design and implementation have a large impact on farm management and overall land use, and therefore on farmland biodiversity (Pe’er et al., 2014). Mandatory set-aside schemes were introduced in the CAP in 1992 and made up a large proportion of all fallows across the EU. All larger cash-crop farms had to set aside (i.e., leave fallow) around 10% of their arable land between 1992 and 2007. In 2007, mandatory set-aside was abolished and, as a result, the total fallow area declined strongly across all EU member states (Tarjuelo et al. 2020, Figure 1). Fallow land area increased across the EU after the 2013 CAP reform, which obliged, from 2015 onwards, most farmers to manage 5% of their arable land as Ecological Focus Areas (EFA) to receive subsidies. Amongst others, the implementation of fallow land was one option for EFAs. However, the area of fallow land did not reach similar levels as during the period of mandatory set-aside (Tarjuelo et al., 2020, Figure 1).

**Figure 1:**
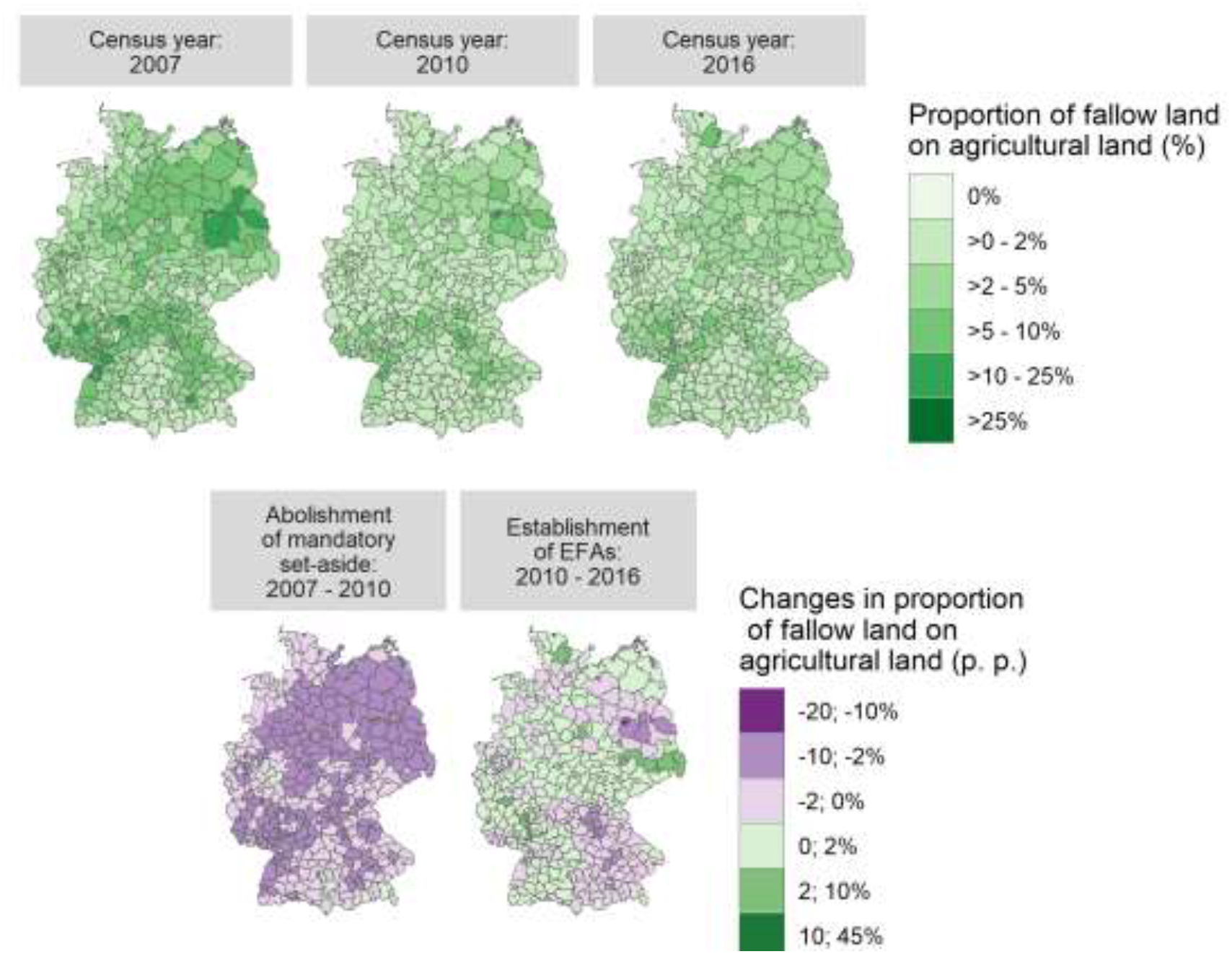
Proportion of fallow land on agricultural land across all German districts at three time steps (upper panel) and changes in the proportion of fallow land in percentage points (p. p.) between 2007 and 2010 (after the abolishment of mandatory set aside in 2007), and between 2010 and 2016 (after the establishment of Ecological Focus Areas) (lower panel).

There is little doubt that farmland biodiversity generally benefits from a high proportion of fallow land in agricultural landscapes (Van Buskirk & Willi, 2004). However, the moderating role of landscape structure on the effectiveness of fallows for promoting farmland biodiversity remains unclear (but see Wretenberg et al. 2010). Landscape complexity can play an important role in shaping biodiversity in agro-ecosystems (Gonthier et al. 2014). Complex agricultural landscapes are composed of an heterogenous mixture of different land-cover types and semi-natural habitats, with a high amount of small woody features (e.g. hedgerows, small woods, scattered trees) (Concepción, Díaz & Baquero, 2008). Increasing the area of fallows in such high-complexity landscapes might have little effect on biodiversity because local species richness might be close to the regional species pool (Tscharntke et al., 2011). In contrast, extremely simplified, “cleared”, agricultural landscapes are dominated by cultivated agricultural land with no or very low amounts of semi-natural habitats and low amount of small woody features. In such low-complexity landscapes, potential source populations of farmland species might be isolated and too depleted to respond to increases in the area of fallows and resources provided (Tscharntke et al., 2012). The ‘intermediate landscape complexity hypothesis’ predicts that local conservation measures will be most effective in promoting species richness in agricultural landscapes of intermediate complexity and less effective in both completely cleared and highly complex landscapes (Tscharntke et al. 2005, 2012). Empirical evidence suggests that species richness does increase in simple landscapes of intermediate complexity containing a low to medium amount of semi-natural habitats, once fallows are provided (Wretenberg et al., 2007, 2010).

The area of fallows in the agricultural landscape also affects the abundance of farmland species. Fallows, compared to productive fields, provide food and undisturbed habitat leading to higher bird population densities and higher reproductive success (Whittingham et al. 2006). It is likely that abundance responses to changes in the area of fallows will be moderated by species-specific ecological requirements, such as the preference of birds to forage and reproduce in open farmland (field-breeder) or in edge habitats (edge-breeder) (Berg & Pärt 1994). Additionally, landscape complexity could also moderate the response of farmland species abundance to changes in the area of fallow land. Fallows were reported to harbor higher abundances of plants compared to field margins especially in simpler landscapes (Ma & Herzon 2014) and the abundance of open farmland bird species in fallows was highest on long-term fallow in simpler landscapes (Toivonen et al 2015).

Here, we investigate the relationships between fallow area and farmland biodiversity spanning three CAP funding periods between 2007 and 2016. We used farmland birds as model organisms, as they are established biodiversity indicators (Gregory et al., 2005), subject to a long history of monitoring (Brlík et al., 2021) and have been suggested to strongly respond to changes in fallow land area (Traba & Morales 2019). In addition, we tested whether the strength of the relationships between species richness and abundance of farmland birds and fallow land area were moderated by species habitat preferences and landscape configurational complexity quantified as the total length of edges between agricultural fields and woody features in the landscape. According to our expectations, we phrased the following hypotheses:

1. Farmland bird species richness and abundance is generally positively related to fallow area, because these serve as high quality habitats allowing high breeding success and survival rates of farmland bird populations.
2. The association of farmland bird abundance with fallow area is moderated by species’ breeding habitat preference: There will be a stronger association of field-breeder and edge-breeder abundance with the proportion of fallows in the landscape compared to that of foraging visitors, because the latter have weaker dependency on fallows and are additionally constrained by the availability of nesting sites such as hedgerows and buildings.
3. The relationship of farmland bird species richness with fallow area is strongest in landscapes of intermediate configurational complexity and weaker in landscapes with low (“cleared”) or high (“complex”) configurational complexity (Tscharntke et al. 2012), because cleared as well as complex landscapes are avoided by several farmland species.

## 2. Materials and Methods

### 2.1. Bird data

We used data on 24 farmland bird species (Table S1) from the German Common Breeding Bird Survey (CBBS) (Kamp et al. 2021). In this monitoring program, more than 2600 sample plots of 1 km^2^ were selected in a randomly stratified way. Up to 1800 of these are surveyed annually (Kamp et al. 2021). Within each plot, experienced volunteer observers walk a predefined route of ca. 3 km length and record all detectable (i.e. optical & acoustical) individuals of all common bird species (without distance limits) and their behavior. Four repeated surveys are conducted in the period between 10 March– 20 June. Standard territory mapping (Bibby et al. 2000) is used to combine the observations into territories along the route. We used the annual number of territories per sample plot and species as a measure of abundance. Plot-level species richness was calculated from the raw data. We classified the selected farmland bird species into three categories based on their ecological requirements and their preference to use arable fields and fallows for breeding or foraging using species-specific habitat associations reported in the literature (Table S1). Species that are known to breed mainly inside fields, including fallows, were categorized as ‘field-breeders’ (n=5), species that prefer the edges of fields and fallows were categorized as ‘edge-breeders’ (n=7). Species that are known to breed outside of fields and fallows (e.g. in hedgerows), but visit those regularly for foraging were classified as ‘foraging visitors’ (n=12). We calculated species richness across all categories and total abundance for each category separately. To account for the fact that some species reach larger population sizes and densities than others, we scaled bird species abundance to values between 0 and 100, 100 corresponding to the maximum abundance reported for the respective species in any year. The species-level scaled abundances were then summed to the nearest integer per category.

To match the bird data to the agricultural census data (see below), data from the years 2007, 2010 and 2016 were used in the analysis. Further, only CBBS plots with at least 10% agricultural area (cropland and managed grasslands) were kept. This resulted in 613, 742 and 948 plots available for analysis for the years 2007, 2010 and 2016. Each CBBS plot was assigned to a German district (NUTS III level) based on its midpoint.

### 2.2. Agricultural census data

To estimate the area of fallow land, we used agricultural census data at the district level across Germany (n=401 districts; Fig. 1). Data on agricultural management are collected in Germany by the regional statistical authorities at the level of individual farms. This includes e.g. the total agricultural area and the area left fallow (or ‘set aside’) per farm. The latest data are from the years 2007, 2010 and 2016. Due to confidentiality restrictions the officially published data have gaps. Therefore, we used a dataset with gaps filled (Gocht & Röder 2014), provided by Thünen Atlas (2021). Given that the districts differ in size, we used the proportion of fallow land in further analysis computed as the fallow land area divided by the cultivated agricultural area. The census years cover three funding periods of the CAP: the ‘set-aside period’ (2007; fallow land area compulsory), the abolition of mandatory set aside (2010; decrease of fallow land area) and the introduction of Ecological Focus Areas as greening measure (2016; increase of fallow land area; Figure 1).

The term “fallow” comprises a wide range of management options that vary widely in terms of vegetation structure and composition (Underwood & Tucker, 2006). In the agricultural census data, fallow land is defined as agricultural land on which there is no production. Fallow land may either be bare land with crop stubbles ploughed in and natural regeneration of vegetation, or sown with plant mixes ranging from simple grass mixtures to limit the establishment of weeds to seed mixtures of wildflowers designed to benefit farmland biodiversity (Nitsch et al., 2016). Fallows can be present for one growing season (rotational or annual fallow) or over several growing seasons (perennial fallow). Fallows can further differ with respect to their size and management, but have in common that they are not fertilized and treated with pesticides. To comply with the CAP’s “minimum maintenance” requirement, the vegetation must be mulched or mown once a year (see Text S2). The agricultural census data used here did not contain any information on fallow management, thus our analysis lumps together a wide variety of fallows with regard to vegetation structure and composition.

### 2.3. Landscape complexity

We quantified landscape complexity within a radius of 1 km around the midpoints of the CBBS plots (“landscape sector”), because farmland birds respond strongly to landscape complexity at this spatial scale (e.g. Winquist et al. 2011) (Figure S1). Landscape complexity was measured in terms of configuration by calculating the total length of edges between woody features and agricultural land for each landscape sector. Spatial information on linear (e.g. hedgerows), patchy (e.g. isolated patches of trees or shrubs) and additional small woody features were gathered from a 2015 pan-European high-resolution remote sensing product (Copernicus Land Monitoring Service 2015). Information on the land-cover types agricultural land (comprising cropland and managed grasslands) and forest (closed deciduous and coniferous forests) were obtained from vector-based high-resolution (1:25000) land-cover data (ATKIS Base DLM, Bundesamt für Kartographie und Geodäsie). We merged the geometries of the woody features from remote sensing with the ATKIS forest geometries and calculated the edge length between these woody geometries and agricultural fields. We subsequently computed the edge density in meter per hectare. Thus, landscapes with high edge density values indicated high configurational complexity and vice versa (see Figure S1). This configurational metric was used instead of a compositional descriptor because woody features such as hedgerows or forest patches are effectively protected under German law since at least 2005 (e.g. § 9 BundesWaldGesetz). We therefore assume constant edge density between 2007 and 2016. Descriptive statistics of the computed variable are given in Table S2.

### 2.4 Data analysis

We modelled species richness, the total scaled bird abundance for the three categories (field-breeders, edge-breeders and foraging visitors) and bird abundance of the 24 species as a function of the covariates.

Given the hierarchical structure of the data, CBBS plots nested in districts, we applied a multilevel modelling approach using a Bayesian framework. This framework enables flexible model building adapted to the hypotheses and the data (Gelman & Hill 2006) and also allows to consider interregional variability. Furthermore, given the hypotheses formulated in the introduction we did not run (sequential) model selection but rather fitted models that were set up according to the hypotheses. This was complemented by a set of residual checks to ensure that inference drawn from the fitted models can be interpreted with sufficient confidence. We included two main covariates, both z-standardized, in the models: proportion of fallow land (% fallow) at the district level and edge density (meter per hectare) in the landscape sector. We also included the agricultural land cover in hectares in the landscape sector to account for differences in habitat availability. Exploration of the data suggested that a square root transformation of fallow proportion improved the linearity of the relationship with the response variables; we therefore transformed the covariate accordingly.

To separate the impacts of the spatial and the temporal gradient of fallow land area on farmland birds (see Oedekoven et al. 2017) we fitted separate models for each of the three years.

The model fitted to the species richness data was of the form:

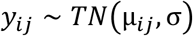

Where *y* is the species richness being modelled based on a truncated normal distribution (*TN*) with a lower bound set at 0 and with expected value *μ* and standard deviation *σ*. Other statistical distributions were explored such as the negative binomial distribution or gaussian distribution with a log link, but the model results showed poorer fit for these distributions compared to the truncated normal distribution with an identity link. The indices are *i*: the individual observations and *j*: the district. The expected values were modelled as follows:

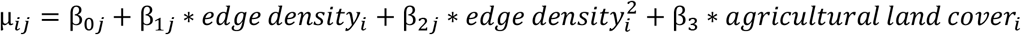

The coefficients β_0_, β_1_ and β_2_ were modelled as follows:

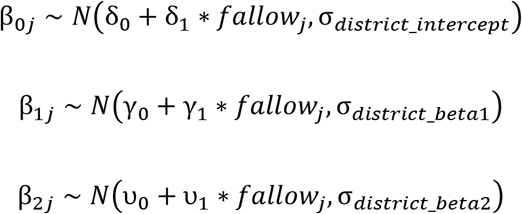

The *σ’s* are independent deviation parameters. The coefficients β_0_, β_1_ and β_2_ were varying between the district in relation to group-level predictors, which is the amount of fallow land reported in the district. The models fitted to the abundance data were of the form:

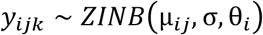

Where *y* corresponds to the scaled abundance data being modelled based on a zero-inflated negative binomial (*ZINB*) distribution with the expected value: μ, the deviation σ and the zero-inflation term θ. The expected values were modelled as follow:

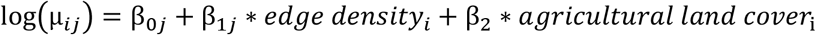

With a similar hierarchical structure as for the species richness model. The coefficient θ was modelled as follow:

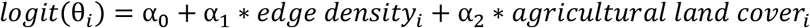

The models were fitted in R (Version 3.6.1) using the brms package v2.16 (Brückner 2017) with weakly informative prior distributions (see Table S3). Default sampling settings were used except for the parameter adapt_delta and max_treedepth which were set to 0.9 and 25, respectively. Posterior distributions were estimated by running four independent chains for 2000 iterations, half of which were used as burn-in for the sampler and discarded. We conducted the following model checks:

- convergence checks ensuring that the Rhat values for all parameters was below 1.1.
- efficiency checks ensuring that the ratio of effective sample size over the number of posterior samples was above 10%.
- posterior predictive checks comparing the density of the observed data against the density of simulated data using the function pp_check from the R package brms.
- spatial autocorrelation of scaled residuals computed using the DHARMa package v0.4 (Hartig 2019) and the posterior_predict function from the brms package. Two aspects of the spatial autocorrelation were checked: (i) the global Moran’s I value derived from the function testSpatialAutocorrelation and (ii) a spline fit to the correlogram of the residuals derived from the function spline.correlog of the package ncf v1.2 (Ottar 2020).

R-square values of the models were computed using the function bayes_R2 from the brms package, both considering all parameters in the models (conditional R-square) and restricting the R-square to the model covariates (marginal R-square). A sensitivity analysis was conducted to explore the impact of individual species on the estimated effects of fallows for the different bird categories. Each species was dropped once at a time from the computation of the total scaled abundance and the models were refitted. The posterior distributions of the fallow slope and the interaction between edge density and fallow were then compared between the initial model and the model without the focal species (see Table S4).

To test our first and third hypothesis, we extracted the posterior draws of the estimated effect of fallows and its interaction with landscape configurational complexity from the models fitted to bird species richness (parameters δ, γ and *v* from the model equations above). The effect of fallows was then computed along a gradient of edge density comprising 95% of the observed values and ranging from 5 (2.5% quantile) to 113 m (97.5% quantile) of edges per hectare.

To test our first and second hypothesis, we compared the strength of the effects of fallows on bird population abundance between the three different bird groups using the posterior draws of the δ and γ model coefficients. We compared the posterior draws between the three bird groups and the 24 species.

## 3. Results

All model parameters were efficiently sampled (effective sample size ratio larger than 10%) and converged (Rhat smaller than 1.1). In addition, posterior predictive checks revealed that the distribution of simulated data based on the model parameters’ posterior distributions matched the distributions of the observed data (Figure S2 and S3). Spatial autocorrelation of the model residuals was generally low (Moran’s I 6e^-3^ to 3e^-2^, Figure S4) and correlograms showed no evidence of spatial signal in the residuals (Figure S5). The models explained between 13% (foraging visitors in 2016) and 50% (field-breeders in 2016) of the variance in the data. When considering only the covariates without the hierarchical terms the models explained between 6% (foraging visitors in 2016) and 32% (field-breeders in 2010) of the variance in the data.

Supporting our first hypothesis, the proportion of fallow land was generally positively associated with farmland bird species richness and bird abundance (Figure 2 and 3).

**Figure 2:**
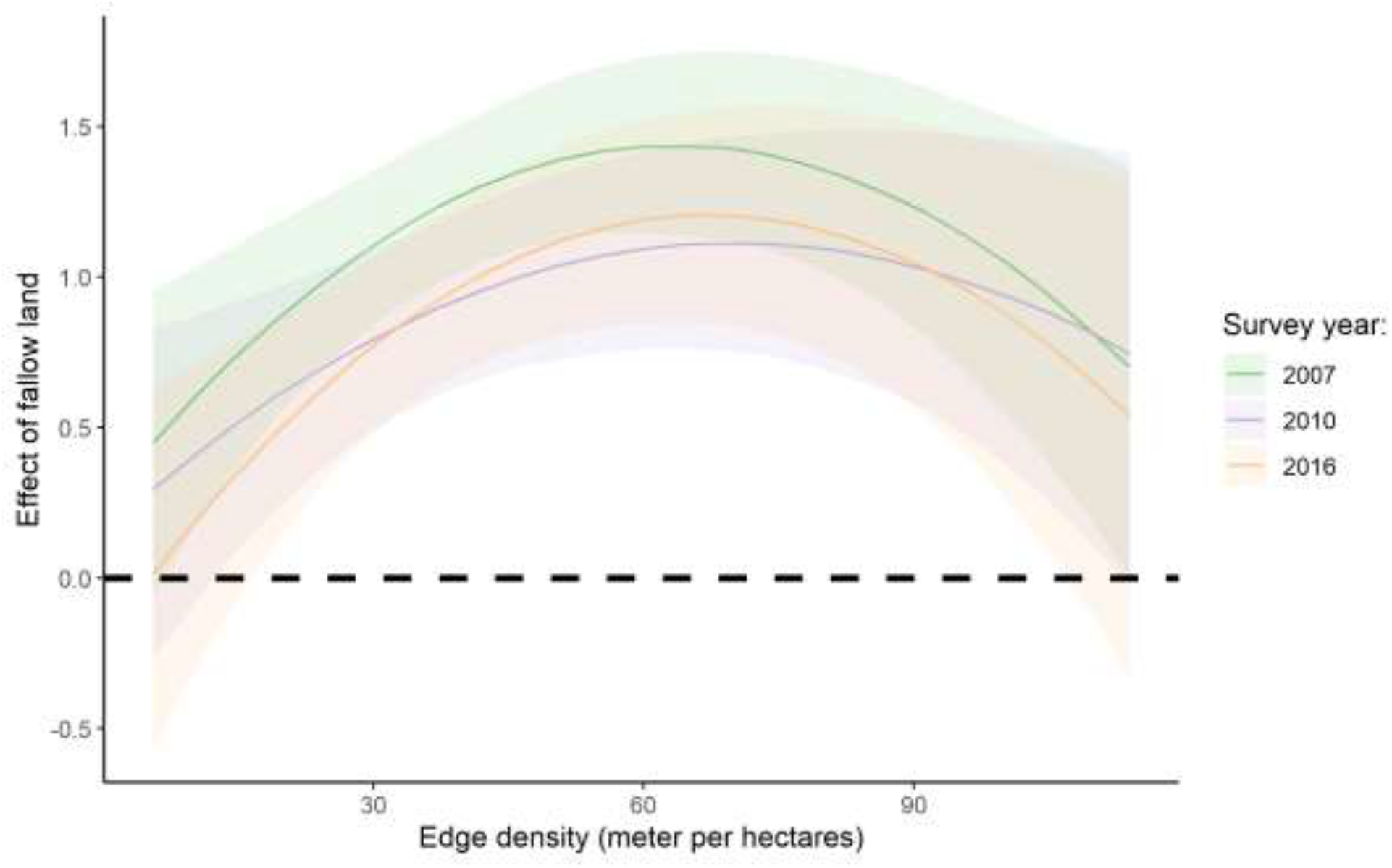
Effect of the proportion of fallow land on farmland bird species richness conditional on landscape configurational complexity (edge density). The thick lines represent the posterior mean of the estimated effect and the contour lines the 95% credible interval around the estimated means.

**Figure 3:**
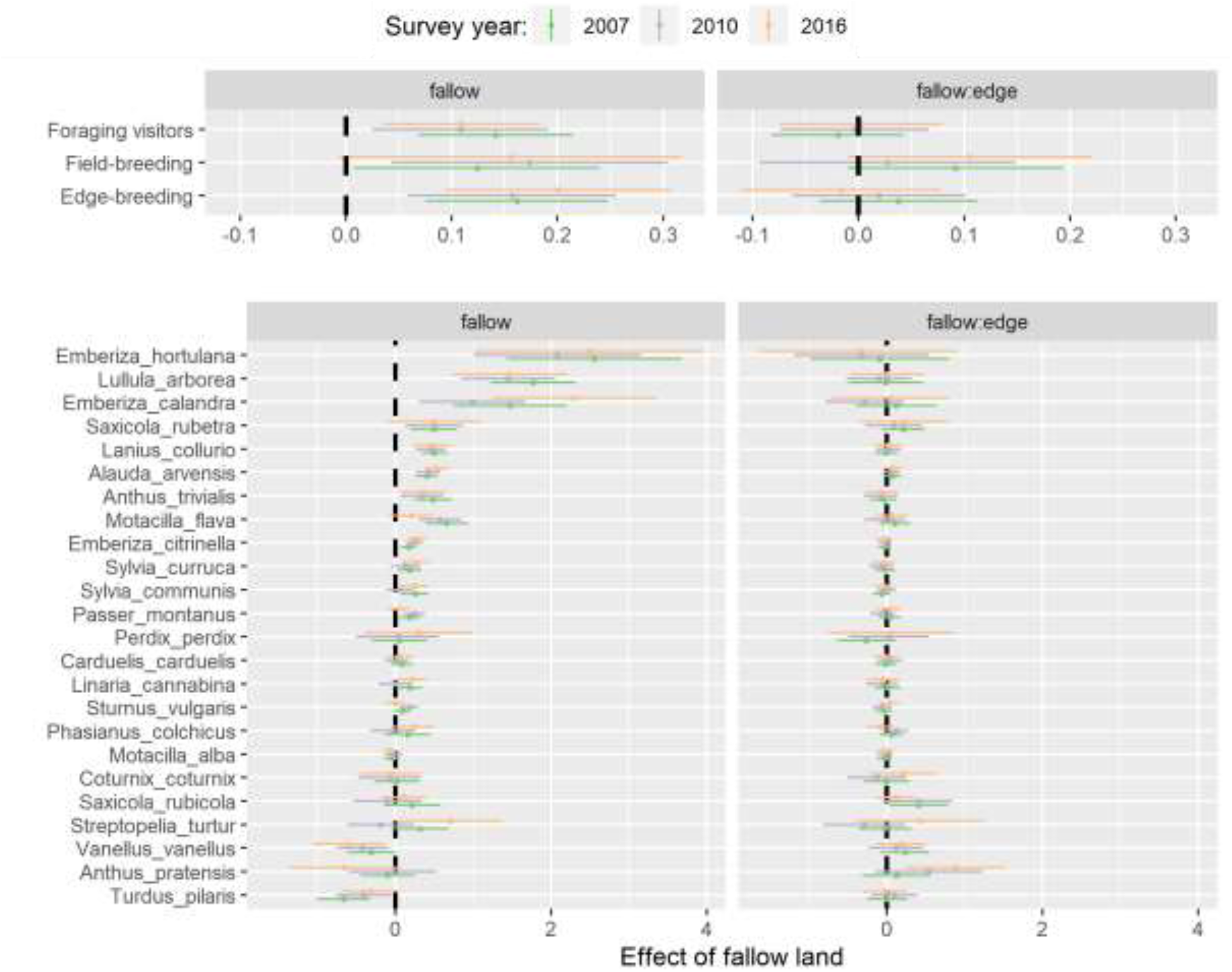
Estimated coefficients (posterior means +-95% CrI) from models relating proportion fallow to bird abundance. The left panels show the effect of the proportion fallow land on the abundance of single species (below) and group-level abundance (above). The right panels show an interacting effect of fallow land and landscape configurational complexity (edge density).

The relationship of species richness with fallows was dependent on landscape configurational complexity and showed a hump-shaped relationship (Figure 2), supporting our third hypothesis. In low-complexity landscapes with edge densities below 14 meter per hectare, the relationship of fallow land with species richness was uncertain in at least one of the focal years (i.e. the 95% credible interval crossed 0). The relationship of fallow land with species richness was strongest in landscapes with around 65 meter of edges per hectare. This represents a high edge density in Germany, as 78% of the landscape sectors had lower edge densities. For landscapes with edge densities higher than 65 meter per hectare the strength of the effect of fallow land declined but at the same time the uncertainty around the estimated effect of fallow land increased markedly, potentially due to the low number of data points at this end of the gradient (Figure 2).

There was strong evidence for a positive association between fallow land and both edge-breeders and foraging visitors with no evidence for a moderating role of landscape complexity on this relationship (Figure 3). Field breeders also showed a tendency towards higher abundances in landscapes with a larger proportion of fallows, but uncertainties were higher. In the survey years 2007 and 2016, there was some evidence of a positive interaction between fallow land and landscape complexity on field-breeders, hinting towards stronger association between fallow land and field-breeders in landscapes with higher edge densities (Figure 3). Comparing the strength of the effect of fallow land between the three species groups in the different years and for different levels of landscapes complexity revealed no clear indication of differences between the species groups with the exception of some evidence towards larger effect of fallow land on field-breeders in landscapes with high edge densities in 2007 and 2016 (Figure S6). At the species-level, there was some evidence for a positive association between fallow land and species-level abundance in at least one of the studied years for a majority of species (Figure 3). Landscape complexity did not appear to mediate these associations for most species and years. Two species (*Vanellus vanellus* and *Turdus pilaris*) showed a negative association with fallow land in at least one of the studied years.

Landscape configurational complexity had a hump-shaped relationship with bird species richness (Figure S7), and positively affected the abundance of both edge-breeders and foraging visitors. An increase of edge density negatively affected the abundance of field-breeders (Figure S8).

## 4. Discussion

We found good support for our first hypothesis stating that farmland bird populations are associated with fallow land area across three CAP funding periods. Consistent with previous albeit more correlative studies at lower data resolution (e.g. Traba & Morales 2019, Busch et al. 2020), our results imply that the loss of fallows in the period 2007 to 2016 is a likely driver of recent farmland bird declines. Our findings indicate that the relationships of farmland bird species richness and abundance with fallow land area were moderated by species’ habitat preferences and landscape complexity, which has remained largely unexplored so far (but see Wretenberg et al 2007). Finally, our study demonstrates that data from a national biodiversity monitoring scheme based on long-term structured surveys can be related to agricultural statistics data to explore associations between land-use and biodiversity, thereby complementing results from targeted smaller-scale field studies.

Comparing the strength of the relationships between fallow land and bird species with varying habitat preferences did not provide support for our second hypothesis. The three species categories (field-breeders, edge-breeders and foraging visitors) showed similar association strength with fallow land. There was, however, some indication for a stronger relationship of fallows with field-breeders compared to foraging visitors in the most complex landscapes. This counterintuitive result could be explained by the fact that the selected CBBS plots showing the highest edge density values in their surroundings were still rather open landscapes (Figure S1) where populations of field-breeders could be maintained, albeit at low densities (Figure S8). In these complex landscapes the abundance of field-breeders is low and particularly at risk from breeding failure due to high predation pressure at forest edges (Ludwig et al 2012). In agricultural landscapes, predators often concentrate near woody features, but predator densities in unproductive features such as sown flower strips are low, especially away from their edges (Laux et al. 2022). Therefore, fallow land might offer safe nesting sites for ground-foraging and ground-nesting birds such as *Alauda arvensis* and *Vanellus vanellus*, while at the same time providing plentiful insect and seed resources (Berg and Pärt 1994).

Providing or retaining fallows will therefore potentially affect a broad range of farmland bird species. Some of these species are amongst the farmland birds that have shown the strongest declines in recent decades such *as Perdix perdix* or *Alauda arvensis* (Kamp et al 2021). Identifying effective conservation measures that have potential to reverse the declines of these species is therefore important. Our results confirm previous studies from other parts of Europe and beyond showing that fallow area and availability might slow down or even reverse negative population trends, e.g. in nine farmland species in Switzerland (Meichtry-Stier et al 2018) and in *Tetrax tetrax* in Spain (Delgado and Moreira 2010).

Our results partially support the ‘intermediate landscape complexity hypothesis’ (Tscharntke et al 2012) postulating that conservation measures will be most effective for restoring high levels of species richness in landscapes with intermediate complexity. Indeed, we found that the effect of an increasing proportion of fallows on farmland bird species richness was lower in landscapes with low edge density compared to landscapes with intermediate edge density. This could be due to the fact that in cleared landscapes (no edges between agricultural land and woody features) the species pool is limited by the lack of woody features required by numerous species and providing fallow land alone is not sufficient to increase farmland bird diversity (Tscharntke et al. 2012). At the other extreme of the gradient in landscape configurational complexity, our results show that the effect of fallow land declines in landscapes with an edge density exceeding 65 meter per hectare. However, for these most complex landscapes the uncertainty around the magnitude of the effect of fallow land increases strongly. This might be due to limitations of the modeling framework and/or the low number of CBBS plots that exhibited a large edge density. However, it seems likely that the effect of fallows declines in these most complex landscapes because farmland bird species richness is already maximized. Increasing the proportion of fallow land would therefore not result in a difference in species richness (Tscharntke et al 2011).

Biodiversity benefits from fallows not only depend on the overall area left non-productive, but also strongly on fallow management (Van Buskrik & Willi 2004, Sanz-Pérez et al. 2021). There is evidence that extensive fallow management targeted to the requirements of individual species can be more effective in increasing local farmland bird abundance than conservation measures that adopt more generic management prescriptions (Sanz-Pérez et al. 2021). For instance, habitat suitability for farmland birds depends on fallow age, i.e. the time since the last cultivation in multi-year fallows. Biodiversity benefits generally increase with fallow age (Staggenborg & Anthes 2021) but declines for fallows left uncultivated for more than ten years have been reported (Lameris et al. 2017).

The submitted German 2023-2027 CAP strategic plan (BMEL 2022) entails the requirement for farmers to leave 4% of their arable land as non-productive features (including e.g. fallows). Based on agricultural census data, in 269 out of 401 (67%) districts the respective proportion of fallow land was below 4% in 2016 (Table 1). Our results imply that increases in non-productive features, such as fallows, to meet this new requirement could lead to increases in farmland bird richness and abundance. However, we speculate that a proportion of 4% fallow land on arable land would not be sufficient to restore, across all districts, the former levels of abundance and species richness in farmland birds that were observed before the strong decrease in fallow land around 2007. Indeed, the districts that would need to increase their proportion of fallows to meet this requirement are not necessarily the ones that showed the most severe losses in fallow land between 2007 and 2016 (Figure 1). Additional CAP instruments based on farmers’ voluntary participation, such as eco-schemes and agri-environmental schemes, will likely lead to additional fallows primarily in less productive regions (cf. Röder and Offermann 2021). These additional fallows will be needed to fulfill the target of 10% of agricultural area covered by high-diversity landscape features set in the EU Biodiversity Strategy to support and restore agricultural biodiversity in Europe (COM 2020).

**Table 1:** Descriptive statistics of percent fallow land (over arable land left of the “/”, and over agricultural land on the right) reported in the agricultural statistics for the years 2007 (mandatory set-aside), 2010 (abolishment of mandatory set-aside) and 2016 (establishment of Ecological Focus Areas). There are 401 districts in Germany.

| Census year | 2007 | 2010 | 2016 |
| --- | --- | --- | --- |
| Minimum – Maximum (%) | 0.00 – 73.10 /<br>0.00 – 14.30 | 0.03 – 35.10 /<br>0.01 – 14.30 | 0.19 – 100 /<br>0.01 – 44.30 |
| Median (%) | 5.72 / 3.83 | 2.33 / 1.55 | 3.08 / 2.01 |
| Number of districts > 4% | 295 / 193 | 113 / 55 | 132 / 58 |
| Number of districts > 10% | 77 / 17 | 17 / 2 | 11 / 2 |

Our study has several limitations. First, the fallow land data extracted from agricultural census data did not contain information on fallow management that drives vegetation structure and composition, which in turn can affect breeding bird species. Future analysis capitalizing on other data sources such as the Integrated Administration and Control System (IACS) could address this limitation (Toth and Kucas 2016). However, the use of such data is subject to privacy laws and access is therefore highly restricted for scientific purposes (Dullinger et al. 2022). Another limitation of our analysis was the fact that fallow land data was solely available at the district level rather than at the level of the CBBS plots. This potentially adds uncertainty and noise to our model estimates especially in large districts where fallow land is not homogeneously distributed across the agricultural area. Future studies using georeferenced information on fallow land such as available in IACS could overcome this limitation. In addition, future evaluations on the impact of fallow land or more broadly of local conservation measures on farmland biodiversity could potentially be improved by combining structured data from national monitoring schemes with opportunistic citizen data to contrast changes before and after the implementation of local conservation measures against population changes of species at the landscape scale (Josefsson et al. 2020).

Based on our results, we make the following recommendations to include in future strategies for fallow land development:

1. Reestablish and maintain a minimal amount of fallow land across all agricultural landscapes, for instance through the enhanced conditionality of the 2023-2027 CAP;
2. Management prescription for fallow land must meet the requirements of the targeted species group and the local context (i.e. landscape complexity). A general key aspect is abandoning farming practice which provoke disturbance during the breeding season of targeted species group;
3. Increase the proportion of fallow land beyond minimal required amount especially: (i) in regions that experienced the strongest loss in fallow land and (ii) in landscapes with an intermediate to high level of configurational complexity, i.e. edge density above 14 m/ha.

In addition to fallow expansion, other non-productive biodiversity-friendly features should be supported by policies to bridge the gap between the 2023–2027 CAP and the EU Biodiversity Strategy targets. These could include restoring species-rich grasslands (Alison et al. 2017) or preserving isolated features such as trees and high-quality hedgerows in agricultural landscapes (Pustkowiak et al. 2021).

## Author contributions

S.K., N.R., J.K. and L.H. conceived the idea; L.H., J.K., C.F., S.K., N.R. and H.B. defined the questions and the methodological approach; L.H., J.K. and C.F. collected the data; L.H. carried out analyses and led the writing; and J.K, C.F., S.K., N.R. and H.B. contributed to the writing.

## Acknowledgements

We especially thank the volunteers of the German Common Breeding Bird Survey (CBBS). We thank Friederike Kunz and Sven Trautmann for help with data management. The German CBBS is coordinated by the Dachverband Deutscher Avifaunisten e. V. and financially supported by the Federal Agency for Nature Conservation through funds provided by the Federal Ministry for the Environment, Nature Conservation and Nuclear Safety. Funding for this research was provided by the German Federal Ministry of Food and Agriculture as part of project “Monitoring der biologischen Vielfalt in Agrarlandschaften”.

