## Supplementary Files for "Losses and gains of fallows impact farmland bird populations over three funding periods of the EU Common Agricultural Policy"

**Contents:**

- Table S1: Classification of the farmland bird species into categories together with species-level prevalence and sample size

- Text S1: References used for the classification of the 24 selected farmland bird species into the categories: field-breeders, edge-breeders and foraging visitors

- Text S2: Management prescription for fallows in Germany.

- Figure S1: Aerial photographs of selected CBBS plots along the landscape structural complexity gradient

- Table S2: Descriptive statistics of the computed landscape configurational complexity (edge density) variable.

- Table S3: Prior distribution of the model coefficients

- Table S4: Results from the sensitivity analysis dropping individual species once at a time from the computation of the scaled abundance and comparing the estimated coefficients to the models fitted with the full species set

- Figure S2: Posterior predictive checks for species richness models

- Figure S3: Posterior predictive checks for category-level abundance models

- Figure S4: Residual spatial autocorrelation

- Figure S5: Correlogram of model residuals

- Figure S6: Comparison of the strength of the effect of fallow land between the different bird categories

- Figure S7: Effect of landscape structural complexity on bird species richness

- Figure S8: Effect of landscape structural complexity on the scaled abundance of field-breeders, edge-breeders and foraging visitors

**Table S1:** Overview of the twenty-four farmland bird species and classification of their primary use of fallow fields. ‘Field breeders’ are species that regularly breed and forage in the center of fallow fields. ‘Edge breeders’ are species breeding on the ground at the edges of fields or nearby structures such as field margins and road verges. ‘Foraging visitors’ are species that breed in non-field structures such as hedgerows, but regularly forage in fallow fields. The classification is based on a review of the literature (Text S1). Prevalence refers to the cumulative number of transects across all three years a species was observed (2007, 2010 and 2016); sample size gives the total number of territories used in the analysis (summed over all three years).

| Species name | English name | Category | Prevalence | Sample size |
| --- | --- | --- | --- | --- |
| *Alauda arvensis* | Eurasian Skylark | Field breeder | 1173 | 14223 |
| *Anthus pratensis* | Meadow Pipit | Field breeder | 249 | 685 |
| *Coturnix coturnix* | Common Quail | Field breeder | 467 | 417 |
| *Motacilla flava* | Western Yellow Wagtail | Field breeder | 570 | 1972 |
| *Vanellus vanellus* | Northern Lapwing | Field breeder | 371 | 930 |
| *Emberiza calandra* | Corn Bunting | Edge breeder | 298 | 821 |
| *Emberiza citrinella* | Yellowhammer | Edge breeder | 1413 | 11354 |
| *Emberiza hortulana* | Ortolan Bunting | Edge breeder | 105 | 280 |
| *Phasianus colchicus* | Common Pheasant | Edge breeder | 725 | 2331 |
| *Perdix perdix* | Grey Partridge | Edge breeder | 236 | 177 |
| *Saxicola rubetra* | Whinchat | Edge breeder | 349 | 528 |
| *Saxicola torquatus* | Common Stonechat | Edge breeder | 396 | 466 |
| *Anthus trivialis* | Tree Pipit | Foraging visitor | 783 | 3018 |
| *Carduelis carduelis* | European Goldfinch | Foraging visitor | 1125 | 2476 |
| *Lanius collurio* | Red-backed Shrike | Foraging visitor | 894 | 1525 |
| *Lullula arborea* | Woodlark | Foraging visitor | 338 | 717 |
| *Linaria cannabina* | Common Linnet | Foraging visitor | 896 | 1922 |
| *Motacilla alba* | White Wagtail | Foraging visitor | 1400 | 3794 |
| *Passer montanus* | Eurasian Tree Sparrow | Foraging visitor | 1108 | 7112 |
| *Streptopelia turtur* | European Turtle-Dove | Foraging visitor | 352 | 387 |
| *Sturnus vulgaris* | Common Starling | Foraging visitor | 1426 | 12289 |
| *Sylvia communis* | Common Whitethroat | Foraging visitor | 1139 | 4074 |
| *Sylvia curruca* | Lesser Whitethroat | Foraging visitor | 1091 | 2054 |
| *Turdus pilaris* | Fieldfare | Foraging visitor | 718 | 1956 |

**Text S1:** References used for the classification of the 24 selected farmland bird species into field breeders, edge breeder and foraging visitors (see Table S1).

Busch, M., Katzenberger, J., Trautmann, S., Gerlach, B., Droeschmeister, R., & Sudfeldt, C. (2020). Drivers of population change in common farmland birds in Germany. *Bird Conservation International*, *30*, 335–354.

Burgess, M. D., Bright, J. A., Morris, A. J., Field, R. H., Grice, P. V., Cooke, A. I., & Peach, W. (2015). Influence of agri-environment scheme options on territory settlement by Yellowhammer (*Emberiza citrinella*) and Corn Bunting (*Emberiza calandra*). *Journal of Ornithology*, *156*, 153–163.

Chamberlain, D., Gough, S., Anderson, G., Macdonald, M., Grice, P., & Vickery, J. (2009). Bird use of cultivated fallow ‘Lapwing plots’ within English agri‐environment schemes. *Bird Study*, *56*, 289–297.

Glutz von Blotzheim, U., Bauer, K., & Bezzel, E. (1966). Handbuch der Vögel Mitteleuropas. Akademische Verlagsgesellschaft

Kirby, W. B., Anderson, G. Q., Grice, P. V., Soanes, L., Thompson, C., & Peach, W. J. (2012). Breeding ecology of Yellow Wagtails *Motacilla flava* in an arable landscape dominated by autumn-sown crops. *Bird Study*, *59*, 383–393.

Kullmann, K., Schneider, R., Fischer, S. (1999). Untersuchungen zur Habitatpräferenz der Grauammer (*Emberiza calandra*) in der Uckermark. *Otis,* 7, 154–160.

Meichtry‐Stier, K. S., Duplain, J., Lanz, M., Lugrin, B., & Birrer, S. (2018). The importance of size, location, and vegetation composition of perennial fallows for farmland birds. *Ecology and Evolution*, *8*, 9270–9281.

Németh, T. M., & Winkler, D. (2017). The impact of unmown refuge-strips on the breeding site fidelity of Common Quail (*Coturnix coturnix*) – a case study. *Magyar Apróvad Közlemények*, *13*, 289–296.

Newton, I. (2004). The recent declines of farmland bird populations in Britain: an appraisal of causal factors and conservation actions. Ibis, 146(4), 579-600.

Schmidt, J. U., Eilers, A., Schimkat, M., Krause-Heiber, J., Timm, A., Siegel, S., Nachtigall, W. & Kleber, A. (2017). Factors influencing the success of within-field AES fallow plots as key sites for the Northern Lapwing *Vanellus vanellus* in an industrialised agricultural landscape of Central Europe. *Journal for Nature Conservation*, *35*, 66–76.

**Text S2**: Prescription for management of fallows in Germany

**General obligations for the management of fallows**

Fertilization and application of pesticides generally prohibited (except with explicit special permits granted by regional environmental or agricultural agencies).

**Additional obligations for fallows receiving CAP support**

- before 2015 (DirektzahlVerpflV):
  - on “normal fallows”: no management between the 01.04. and 30.06.; once a year the vegetation must be mulched or mown and the vegetation removed
  - on fallows in Agri-environmental schemes (AES): special management restrictions according to the scheme apply and overrule general obligations
  - on fallows due to implementation of EU environmental legislation: special management restrictions apply and overrule general obligations
  - where landscape features are defined as fallows (e.g. hedges, groves): must be maintained, no additional management restrictions apply
- after 2015 (AgrarzahlVerpflV):
  - As before but on fallows in ecological focus areas:
    - “Normal year”: no management between 01.04. and 15.07. Outside this period ploughing / harrowing with immediate reseeding is possible
    - Last year before planting a normal crop: no management between 01.04. and 15.07. Ploughing not before 31.07.

Fallows keep the legal status of “arable land” (even when not included in a crop rotation for more than five years) in case they are implemented according to the Greening, AES or comparable schemes.

Fallows can also be established on grassland and permanent crops. However, there is neither a legal advantage of having fallow on grassland and permanent crops, nor a requirement to have fallows on these land uses. In addition, the obligations for minimum maintenance are identical for fallows on grassland and permanent crops to the normal minimum management requirements. Therefore, hardly any farmer leaves grassland and permanent crops fallow.

Main references:

AgrarzahlVerpflV (Verordnung über die Einhaltung von Grundanforderungen und Standards im Rahmen unionsrechtlicher Vorschriften über Agrarzahlungen (Agrarzahlungen-Verpflichtungenverordnung) 2014 Agrarzahlungen-Verpflichtungenverordnung vom 17. Dezember 2014 (BAnz AT 23.12.2014 V1), https://www.gesetze-im-internet.de/agrarzahlverpflv/

DirektzahlVerpflV (Verordnung über die Grundsätze der Erhaltung landwirtschaftlicher Flächen in einem guten landwirtschaftlichen und ökologischen Zustand (Direktzahlungen-Verpflichtungenverordnung DirektZahl-VerpflV)) (2004) Bundesgesetzblatt Jahrgang 2004 Teil I Nr. 58, ausgegeben am 12.11.2004, Seite 2778-2784. https://www.bgbl.de/xaver/bgbl/start.xav#__bgbl__%2F%2F*%5B%40attr_id%3D%27bgbl104s2778.pdf%27%5D__1633267031709

**Figure S1:** Aerial photographs from Google Satellite of ‘landscape sectors’ around selected CBBS plot locations along the landscape configurational complexity gradient, characterized by the total length of edges between woody features and agricultural land. The blue circles represent the 1km buffer around the centroids of the CBBS plots.

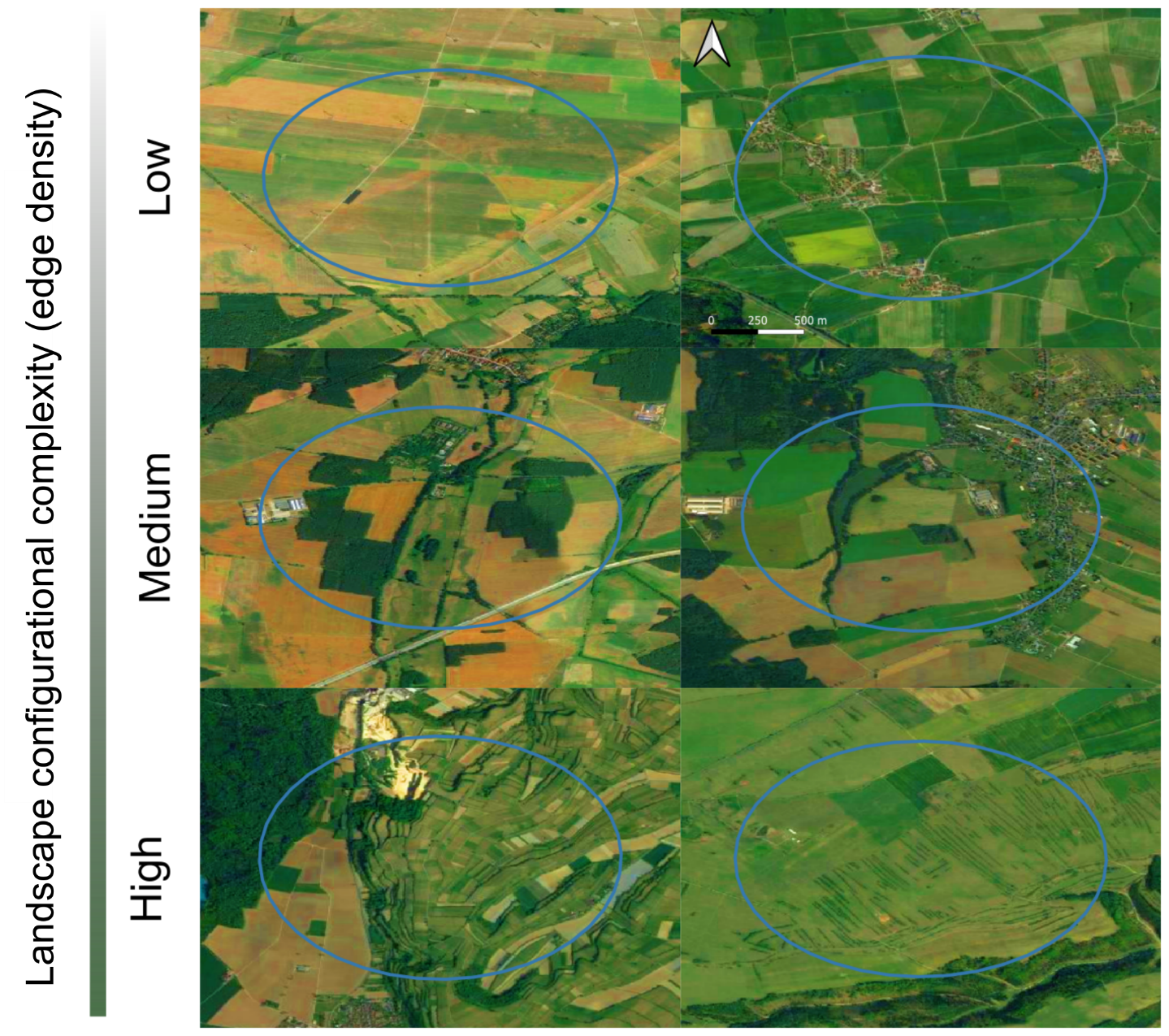

**Table S2**: Descriptive statistics of the landscape configurational complexity (edge density) variable.

|  | Minimum | 25% quantile | Median | Mean | 75% quantile | Maximum |
| --- | --- | --- | --- | --- | --- | --- |
| Edge density (meter per hectare) | 0.00 | 27.22 | 41.91 | 47.01 | 62.85 | 229.90 |

**Table S3:** Prior distributions of the model coefficients, see the model equation in the main text for definitions of the different coefficients

| Coefficient | Distribution | Distribution parameters |
| --- | --- | --- |
| β | Normal | Mean = 0, sd = 2.5 |
| σ (species richness) | Student’s t | Df = 3, mean = 0, sd = 2.5 |
| σ (scaled abundance) | Gamma | Shape = 0.01, scale = 0.01 |
| σ (random effect variation) | Student’s t | Df = 3, mean = 0, sd = 2.5 |

**Table S4:** Results of the sensitivity analysis dropping individual species once at a time from the computation of the scaled abundance and comparing the estimated coefficients to the models fitted with the full species set. Reported are the posterior probabilities of the following comparison: coefficient original model < coefficient of sensitivity model. A value of 0 (or 1) indicates strong evidence that dropping the focal species led to smaller (or larger) model coefficients.

| Category | Species | Year | Slope fallow | Fallow:edge interaction |
| --- | --- | --- | --- | --- |
| Edge-breeders | *Emberiza calandra* | 2007 | 0,574 | 0,5995 |
| Edge-breeders | *Emberiza calandra* | 2010 | 0,59725 | 0,56075 |
| Edge-breeders | *Emberiza calandra* | 2016 | 0,59125 | 0,51275 |
| Edge-breeders | *Emberiza citrinella* | 2007 | 0 | 0,3755 |
| Edge-breeders | *Emberiza citrinella* | 2010 | 0 | 0,6045 |
| Edge-breeders | *Emberiza citrinella* | 2016 | 0,0005 | 0,60375 |
| Edge-breeders | *Emberiza hortulana* | 2007 | 0,65125 | 0,50025 |
| Edge-breeders | *Emberiza hortulana* | 2010 | 0,51125 | 0,3795 |
| Edge-breeders | *Emberiza hortulana* | 2016 | 0,6885 | 0,56 |
| Edge-breeders | *Perdix perdix* | 2007 | 0,0485 | 0,7285 |
| Edge-breeders | *Perdix perdix* | 2010 | 0,21575 | 0,64775 |
| Edge-breeders | *Perdix perdix* | 2016 | 0,1225 | 0,56675 |
| Edge-breeders | *Phasianus colchicus* | 2007 | 0,151 | 0,65925 |
| Edge-breeders | *Phasianus colchicus* | 2010 | 0,58425 | 0,589 |
| Edge-breeders | *Phasianus colchicus* | 2016 | 0,35725 | 0,847 |
| Edge-breeders | *Saxicola rubetra* | 2007 | 0,051 | 0,55375 |
| Edge-breeders | *Saxicola rubetra* | 2010 | 0,14975 | 0,6595 |
| Edge-breeders | *Saxicola rubetra* | 2016 | 0,11225 | 0,5945 |
| Edge-breeders | *Saxicola rubicola* | 2007 | 0,09225 | 0,403 |
| Edge-breeders | *Saxicola rubicola* | 2010 | 0,27075 | 0,483 |
| Edge-breeders | *Saxicola rubicola* | 2016 | 0,1615 | 0,466 |
| Field-breeders | *Alauda arvensis* | 2007 | 0,1765 | 0,22125 |
| Field-breeders | *Alauda arvensis* | 2010 | 0,1875 | 0,2295 |
| Field-breeders | *Alauda arvensis* | 2016 | 0,03825 | 0,57175 |
| Field-breeders | *Anthus pratensis* | 2007 | 0,952 | 0,51675 |
| Field-breeders | *Anthus pratensis* | 2010 | 0,802 | 0,27875 |
| Field-breeders | *Anthus pratensis* | 2016 | 0,963 | 0,18525 |
| Field-breeders | *Coturnix coturnix* | 2007 | 0,947 | 0,5755 |
| Field-breeders | *Coturnix coturnix* | 2010 | 0,764 | 0,44425 |
| Field-breeders | *Coturnix coturnix* | 2016 | 0,8695 | 0,6115 |
| Field-breeders | *Motacilla flava* | 2007 | 0,582 | 0,57525 |
| Field-breeders | *Motacilla flava* | 2010 | 0,51825 | 0,51125 |
| Field-breeders | *Motacilla flava* | 2016 | 0,74925 | 0,633 |
| Field-breeders | *Vanellus vanellus* | 2007 | 0,96525 | 0,82775 |
| Field-breeders | *Vanellus vanellus* | 2010 | 0,979 | 0,74175 |
| Field-breeders | *Vanellus vanellus* | 2016 | 0,9825 | 0,86325 |
| Foraging visitors | *Anthus trivialis* | 2007 | 0,38275 | 0,5025 |
| Foraging visitors | *Anthus trivialis* | 2010 | 0,3745 | 0,461 |
| Foraging visitors | *Anthus trivialis* | 2016 | 0,32225 | 0,5445 |
| Foraging visitors | *Carduelis carduelis* | 2007 | 0,50975 | 0,49225 |
| Foraging visitors | *Carduelis carduelis* | 2010 | 0,53175 | 0,37875 |
| Foraging visitors | *Carduelis carduelis* | 2016 | 0,505 | 0,47975 |
| Foraging visitors | *Lanius collurio* | 2007 | 0,37525 | 0,42225 |
| Foraging visitors | *Lanius collurio* | 2010 | 0,3805 | 0,367 |
| Foraging visitors | *Lanius collurio* | 2016 | 0,41325 | 0,43475 |
| Foraging visitors | *Linaria cannabina* | 2007 | 0,49925 | 0,439 |
| Foraging visitors | *Linaria cannabina* | 2010 | 0,56525 | 0,41975 |
| Foraging visitors | *Linaria cannabina* | 2016 | 0,4405 | 0,51225 |
| Foraging visitors | *Lullula arborea* | 2007 | 0,26725 | 0,4815 |
| Foraging visitors | *Lullula arborea* | 2010 | 0,32575 | 0,481 |
| Foraging visitors | *Lullula arborea* | 2016 | 0,317 | 0,4645 |
| Foraging visitors | *Motacilla alba* | 2007 | 0,70425 | 0,5085 |
| Foraging visitors | *Motacilla alba* | 2010 | 0,73025 | 0,52475 |
| Foraging visitors | *Motacilla alba* | 2016 | 0,7615 | 0,55 |
| Foraging visitors | *Passer montanus* | 2007 | 0,46325 | 0,43675 |
| Foraging visitors | *Passer montanus* | 2010 | 0,385 | 0,486 |
| Foraging visitors | *Passer montanus* | 2016 | 0,592 | 0,38325 |
| Foraging visitors | *Streptopelia turtur* | 2007 | 0,49025 | 0,47125 |
| Foraging visitors | *Streptopelia turtur* | 2010 | 0,519 | 0,43825 |
| Foraging visitors | *Streptopelia turtur* | 2016 | 0,455 | 0,4405 |
| Foraging visitors | *Sturnus vulgaris* | 2007 | 0,47775 | 0,452 |
| Foraging visitors | *Sturnus vulgaris* | 2010 | 0,388 | 0,51225 |
| Foraging visitors | *Sturnus vulgaris* | 2016 | 0,60875 | 0,39525 |
| Foraging visitors | *Sylvia communis* | 2007 | 0,47525 | 0,483 |
| Foraging visitors | *Sylvia communis* | 2010 | 0,585 | 0,324 |
| Foraging visitors | *Sylvia communis* | 2016 | 0,47075 | 0,4955 |
| Foraging visitors | *Sylvia curruca* | 2007 | 0,5065 | 0,42875 |
| Foraging visitors | *Sylvia curruca* | 2010 | 0,51375 | 0,45975 |
| Foraging visitors | *Sylvia curruca* | 2016 | 0,34425 | 0,523 |
| Foraging visitors | *Turdus pilaris* | 2007 | 0,79325 | 0,5225 |
| Foraging visitors | *Turdus pilaris* | 2010 | 0,637 | 0,41625 |
| Foraging visitors | *Turdus pilaris* | 2016 | 0,70525 | 0,4975 |

**Figure S2:** Posterior predictive checks for species richness from left to right for the years 2007, 2010 and 2016. The tick blue lines represent the density distribution of the data used to fit the models and the light blue lines represent 50 distributions simulated from posterior draws of the model parameters.

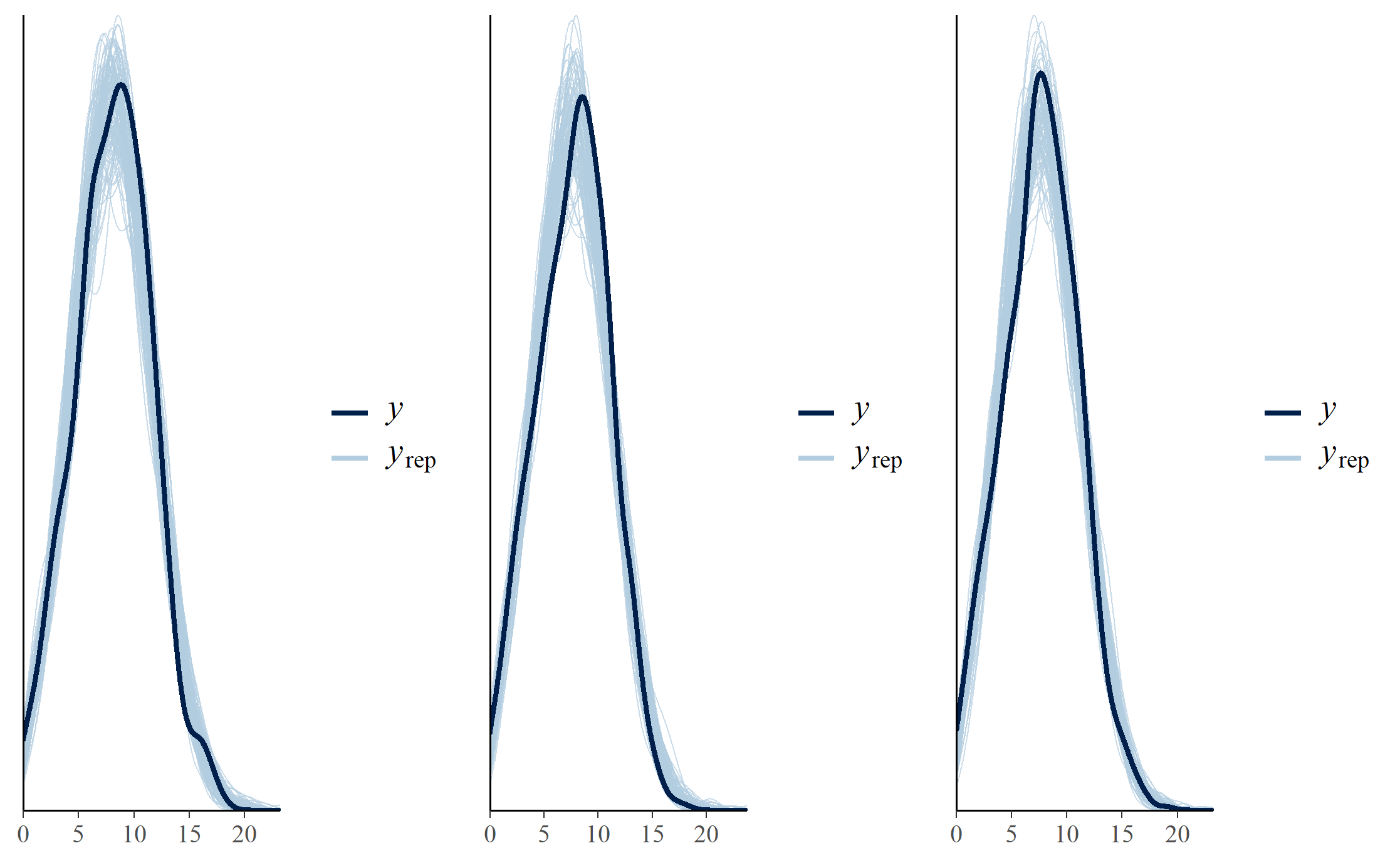

**Figure S3:** Posterior predictive checks for the three species groups, from top to bottom for the edge-, field- and shrub-breeders and from left to right for the years 2007, 2010 and 2016. The tick blue lines represent the density distribution of the data used to fit the models and the light blue lines represent 50 distributions simulated from posterior draws of the model parameters.

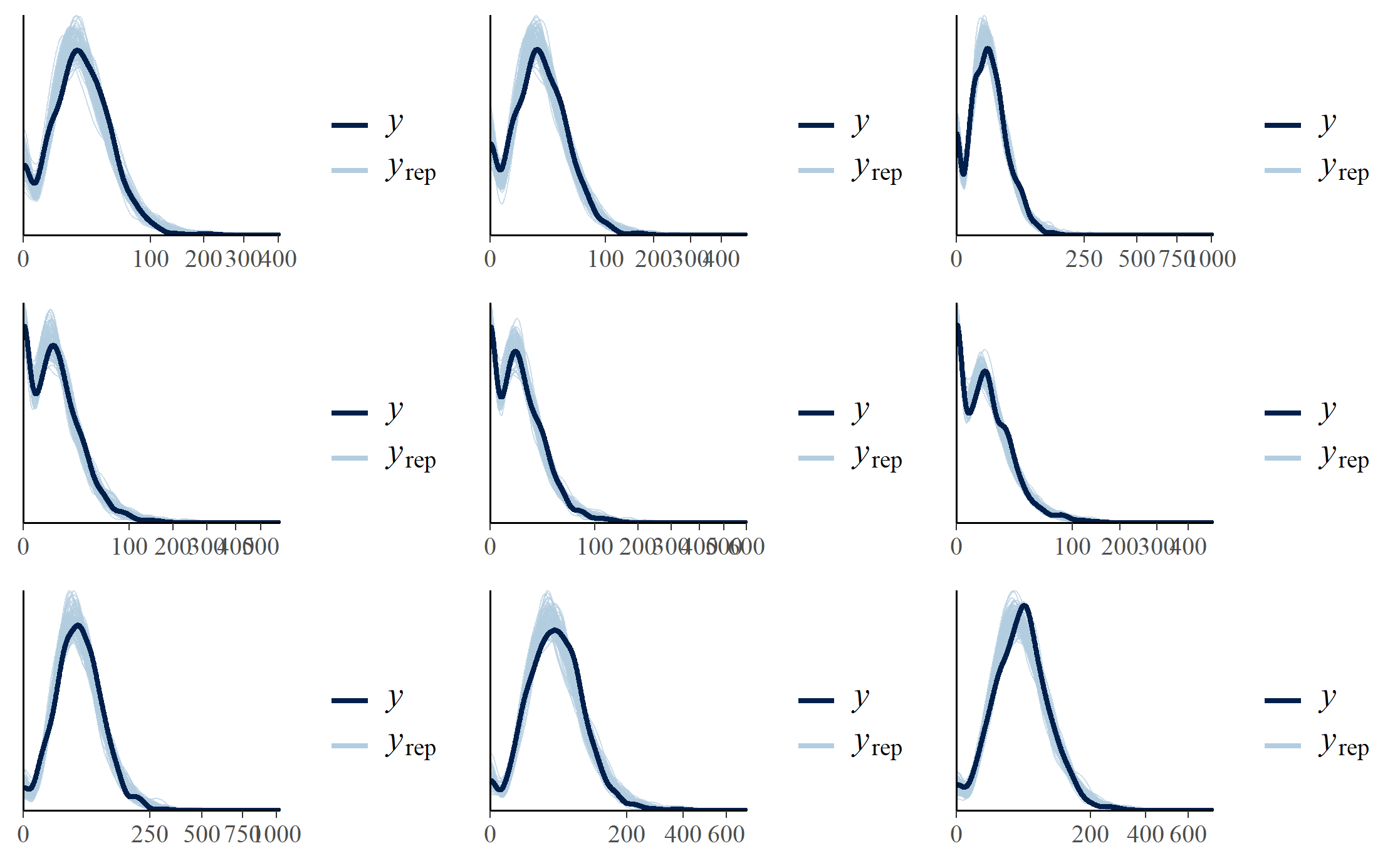

**Figure S4:** Residual spatial autocorrelation with Moran’s I of the species richness and the group-level abundance models. The residuals were computed from the posterior predictive distribution and scaled using the package DHARMa.

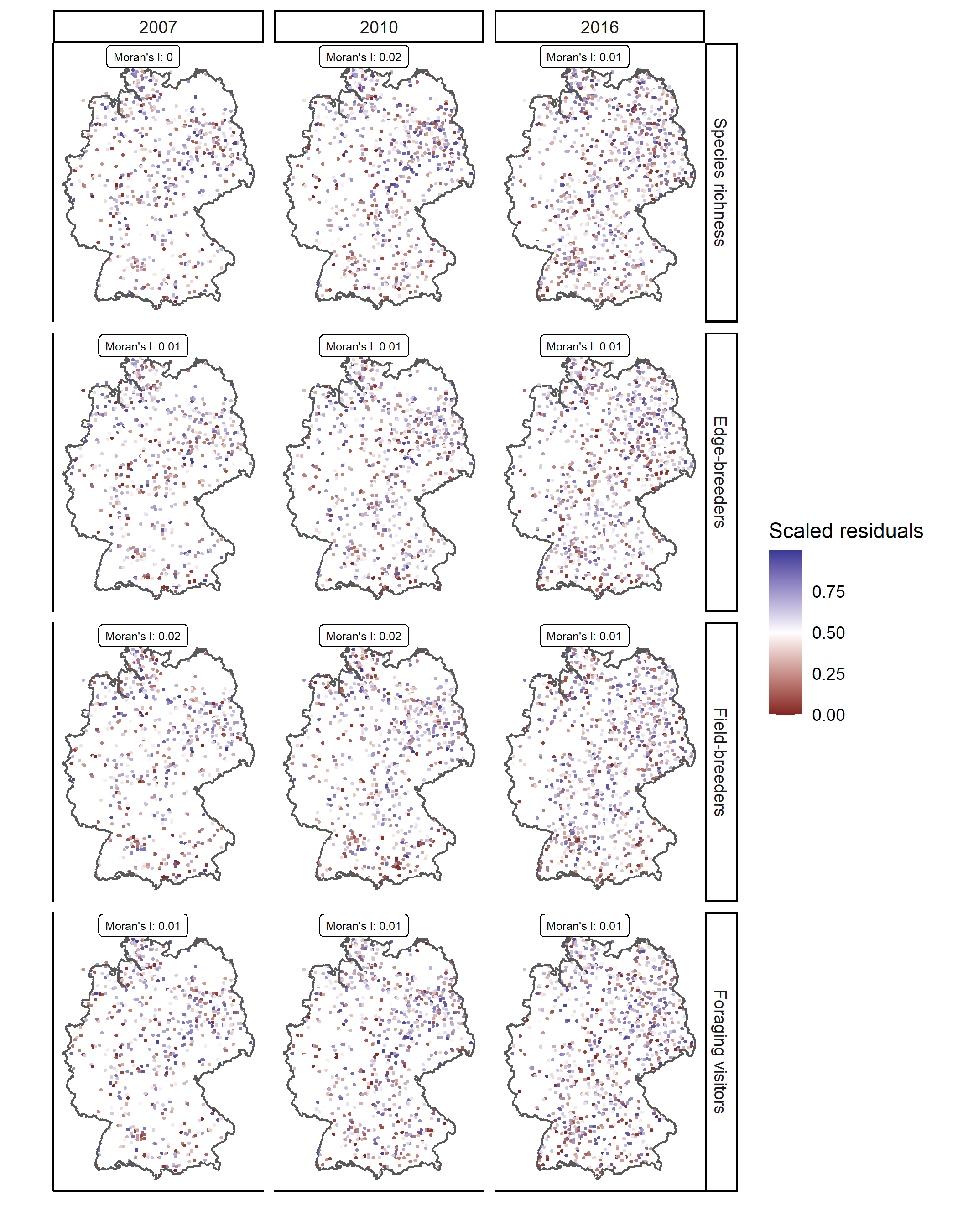

**Figure S5:** Correlogram of the model residuals for the different response variables in the rows and the years in the columns. The residuals were computed from the posterior predictive distribution and scaled using the package DHARMa. A smooth spline was then fitted using the package ncf to the observed relation between correlations of the model scaled residuals and their distance with 1000 bootstraps to derive 95% confidence bands.

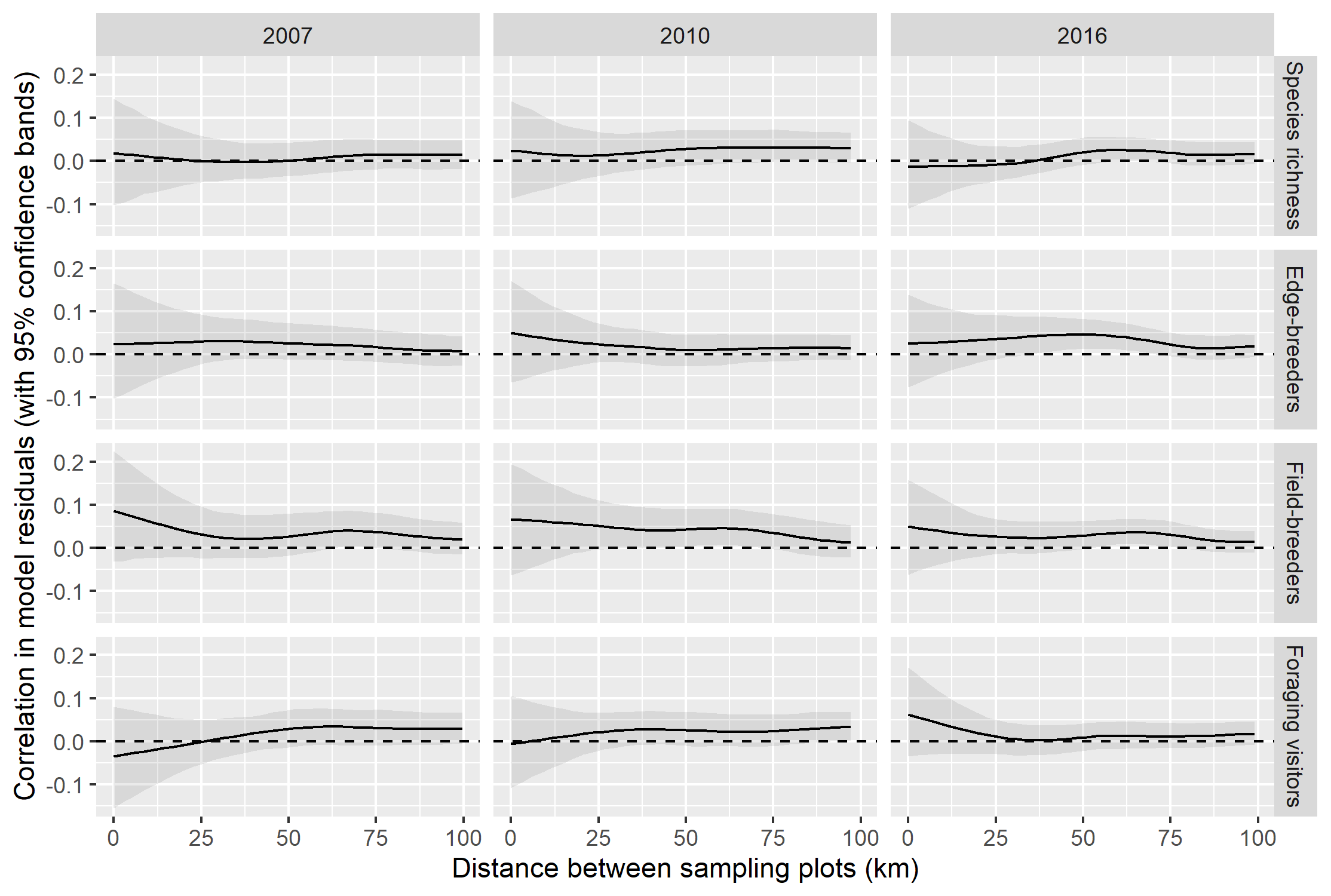

**Figure S6:** Effect of fallow land on group-level bird abundance as predicted from the fitted yearly models. The predictions were derived for three different edge densities: 17 meter per hectare (low edge density, 10% quantile), 47 meter per hectare (medium edge density, median) and 83 meter per hectare (high edge density, 90% quantile). The dots represent the median and the vertical lines the 95% credible intervals.

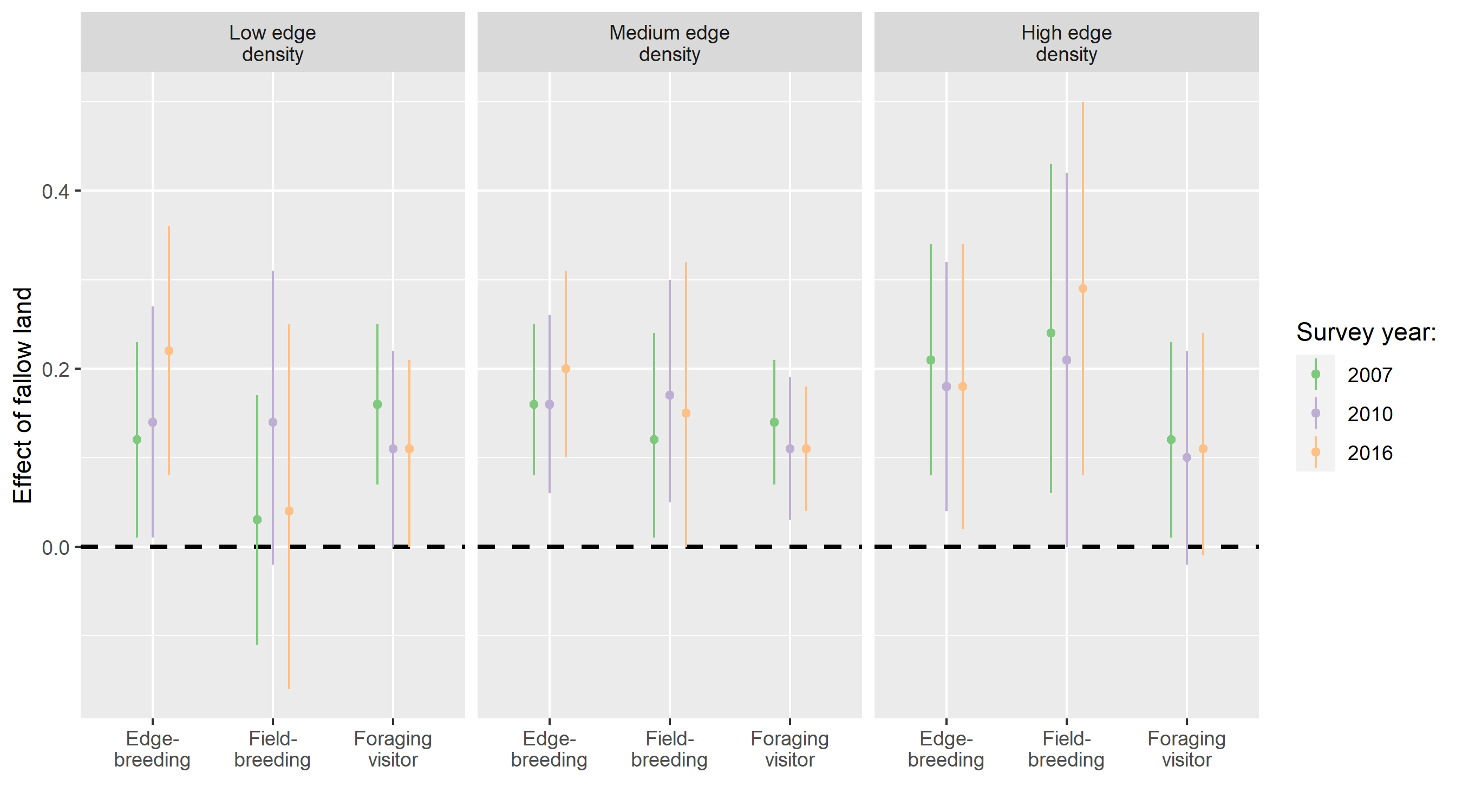

**Figure S7:** Effect of landscape configurational complexity (edge density) on bird species richness as predicted from the fitted yearly models. The predictions were derived for three different proportions of fallow land: 0, 4 and 10%. The lines represent the median and the confidence bands the 95% credible intervals.

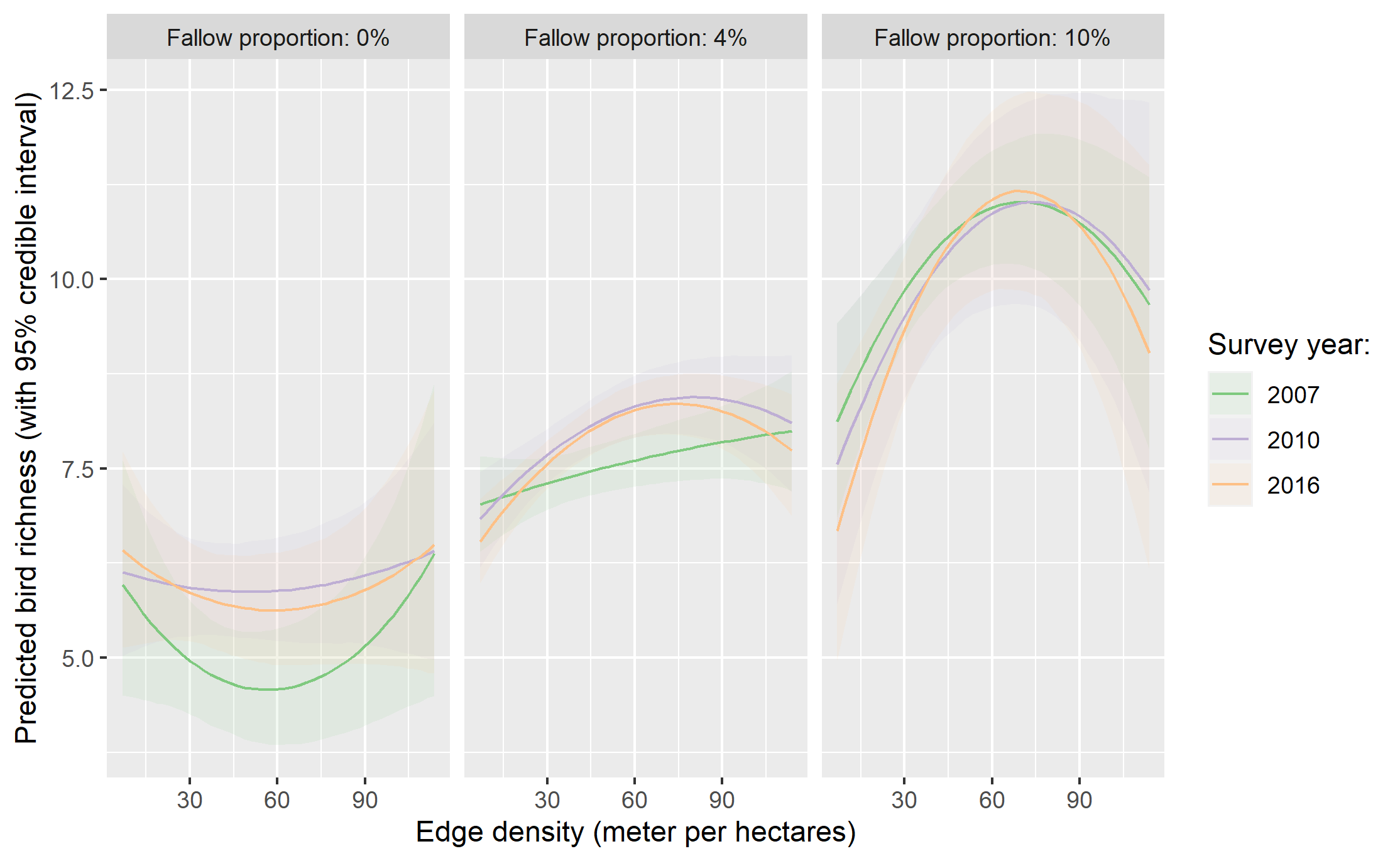

**Figure S8:** Effect of landscape configurational complexity (edge density) on bird species abundance at the group-level as predicted from the fitted yearly models. The predictions were derived for three different proportions of fallow land: 0, 4 and 10%. The lines represent the median and the confidence bands the 95% credible intervals.
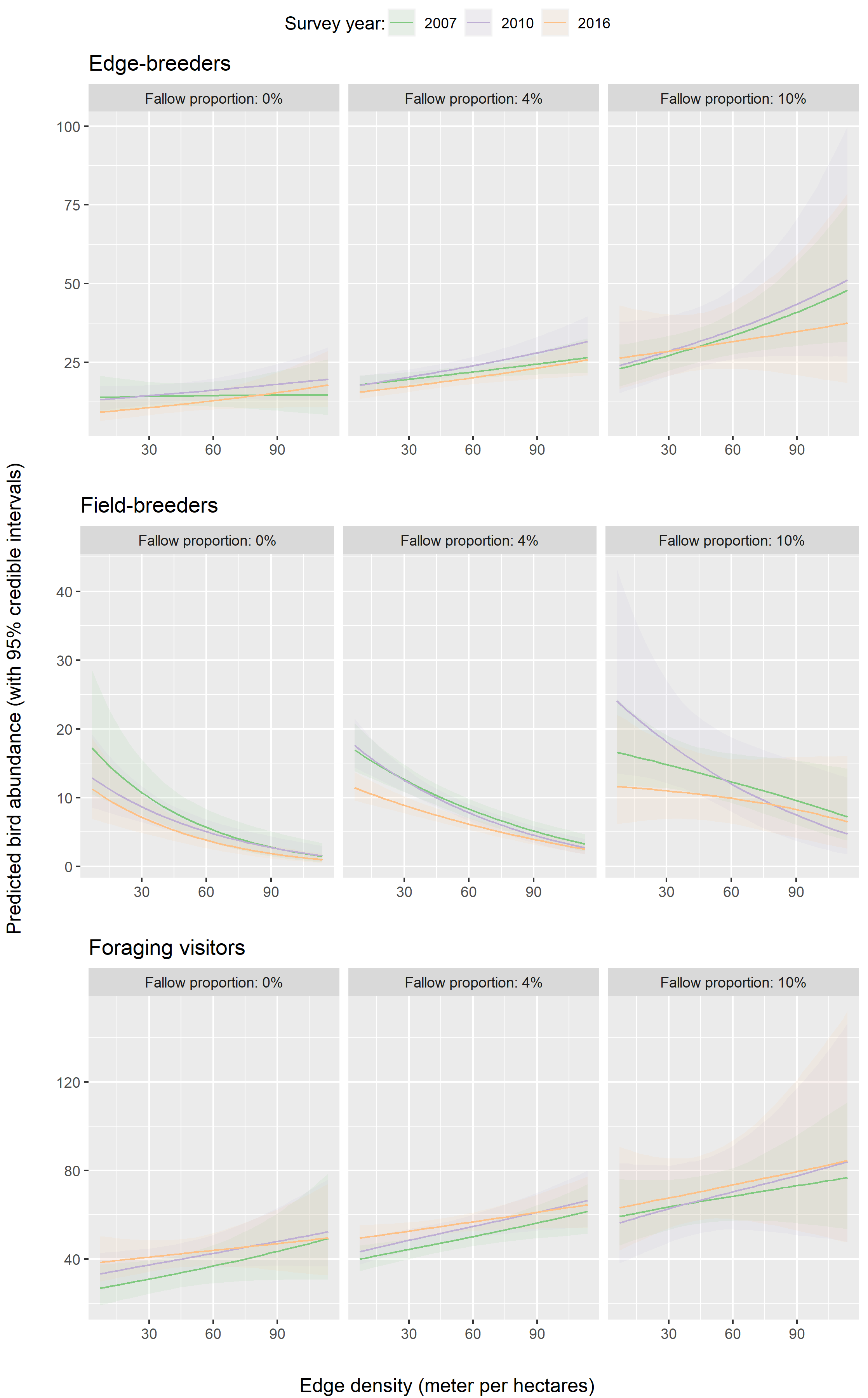
